# Using Virtual Patients to Evaluate Dosing Strategies for Bispecifics with a Bell-Shaped Efficacy Curve

**DOI:** 10.1101/2025.05.20.655186

**Authors:** Jana L. Gevertz, Irina Kareva

**Author notes:** Corresponding author’s.

## Abstract

Bispecific antibodies can be broadly divided into two categories: those that are pharmacologically active as either dimers bound to a single target or as trimers bound to both targets, and those that are only active as trimers. Dose selection of trimer-based bispecifics poses a unique challenge, as toxicity increases with dose, but efficacy does not. Instead, trimer-driven bispecifics have a bell-shaped efficacy curve, for which both under- and over-dosing can cause a decrease in efficacy. To address the challenge of dose selection for trimer-based bispecifics, we develop a semi-mechanistic pharmacokinetic/pharmacodynamic model of one such bispecific, teclistamab. By introducing variability in key patient-specific parameters, we find that the currently selected phase II recommended dose of 1.5 mg/kg administered subcutaneously weekly falls within the calculated optimal range for maximizing concentration of the pharmacologically active trimer for a broad population. We next explore different strategies for patient stratification based on pre-treatment levels of measurable biomarkers. We discover that significantly more variability across subpopulations is predicted when the drug is administered every two weeks as compared to weekly administration, and that higher doses generally result in more interpatient variability. Further, the pharmacologically active trimers are predicted to be maximized at different doses for different subpopulations. These findings underscore the potential for model-supported patient stratification based on measurable biomarkers, offering a middle ground between population-level approaches and fully personalized medicine.

## Introduction

A bispecific antibody is a type of antibody that can simultaneously bind two different antigens or epitopes. Since the approval of the first bispecific antibody catumaxomab in 2009 (which has since been withdrawn), over a dozen bispecifics have been approved (1). Bispecific antibodies can vary in their mechanism of action, with some exhibiting efficacy if bound to a single target, forming dimers (drug + target complexes) and others requiring simultaneous binding to both targets, forming pharmacologically active trimers (drug + target 1 + target 2). If both dimers and trimers contribute to the drug’s efficacy, then the drug effectively acts as a combination of two co-administered monotherapies, with the advantage of being able to co-localize targeting in time and space (i.e., in the same site of action at the same time), and without the need to monitor two separate safety profiles. For instance, cadonilimab targets both PD-1 and CTLA-4 receptors (2), and engagement with either target can be efficacious. Similarly, ivonescimab, which binds both PD-1 and the vascular endothelial growth factor VEGF (3), can have efficacy when bound to either of the arms.

Most of the approved bispecifics are used for targeted cancer therapies, acting to create a direct link between cancer cells and immune cells. These types of bispecifics are referred to a T cell engagers (TCEs), and they typically bind the CD3 receptor on immune cells on one arm and a tumor-specific antigen on the other (4). These drugs can have remarkable efficacy, as seen with glofitamab (brand name Columvi) and epcoritamab (brand name Epkinly) in the treatment of diffuse large B cell lymphoma. Both target CD20 on B cells and CD3 on T cells and have revolutionized outcomes for their respective indications since their approval in 2023 (5,6). While often highly efficacious, these drugs can have serious side effects, most notably cytokine release syndrome, a dangerous immune response characterized by hypersecretion of pro-inflammatory cytokines that can lead to immune hyperactivation and induce various degrees of organ damage (7–9). Therefore, selecting doses that balance efficacy with toxicity is particularly important for trimer-based TCEs.

This task is complicated by the fact that the efficacy curve for such trimer-based drugs is “bell-shaped”. Not only do such drugs have the standard problem of insufficient target engagement and therefore suboptimal efficacy at low doses, but high doses can be equally problematic. Overdosing can result in a stoichiometric shift away from the pharmacologically active trimer (PAT) towards non-efficacious dimers (10,11). This property contrasts with a more typical dose-efficacy Emax curve, where efficacy is expected to plateau rather than decrease with higher concentrations. That is, when both dimers and trimers contribute to efficacy, increasing drug concentration primarily increases target engagement, leading to a classic Emax-shaped dose-response (examples would include the aforementioned cadonolimab and ivonescimab). However, when only trimers drive efficacy, excess drug can saturate individual targets separately, favoring non-productive dimers over pharmacologically active trimers and resulting in a bell-shaped dose-efficacy relationship (such as for epcoritamab, glofitamab or teclistamab).

Thus, for bispecifics in which both dimers and trimers contribute to efficacy, while increasing the dose poses toxicity concerns, it is not expected to result in a decrease in efficacy. However, for trimer-based bispecifics with a bell-shaped efficacy curve, overdosing can lead both to increased toxicity and a loss of efficacy. For these types of drugs, selecting an appropriate dose for most patients is both critically important and challenging.

To explore computational approaches to address this issue, as a test example, we focus on teclistamab (brand name Tecvayli), a bispecific antibody with trimer-based efficacy currently used in refractory or relapsed multiple myeloma (12). On one arm, this bispecific targets CD3 on T cells. On the other arm, it targets the B cell maturation antigen (BCMA), a protein that is often over-expressed on multiple myeloma (MM) cells in the bone marrow. Teclistamab is expected to be efficacious when it is bound to both targets; it is not expected to have efficacy as a dimer, i.e., when bound to only one target. Therefore, estimating the dose that is expected to maximize the drug’s efficacy requires an estimation of the concentration of the pharmacologically active trimer in the tumor microenvironment (TME). Interpatient variability in target expression, distribution kinetics, and target turnover have the potential to influence the pharmacological response curve.

In this manuscript, we introduce a semi-mechanistic pharmacokinetic/pharmacodynamic (PK/PD) model of teclistamab. We use the model to describe the dynamics of the drug and its interaction with its targets to estimate the concentration of the pharmacologically active trimer in the TME and show how this leads to the expected bell-shaped efficacy curve. We next introduce variability in parameters that most sensitively impact the formation of trimers in the TME to generate a cohort of virtual patients. Our model rediscovers that the phase II recommended dose of 1.5 mg/kg administered subcutaneously weekly falls within the optimal range for the average patient in the virtual population. We then leverage virtual population analysis by benchmarking against clinically measurable characteristics. Specifically, we stratify the virtual patients into four subgroups, and we study how the optimal protocol is impacted by these patient characteristics. We conclude with a discussion of the possibility of model-guided semi-personalized therapy as a potential middle ground between allowing the “average” patient to drive dose selection, and fully personalized therapy.

## Methods

We propose a semi-mechanistic model that describes the dynamics of teclistamab as it interacts with its targets in the plasma and the bone marrow: CD3, soluble BCMA and membrane-bound BCMA. The overall objective of the analysis is to maximize the pharmacologically active trimer (PAT = Drug + CD3 + mBCMA). A full description of the model can be found in Supplementary Information. The schematic structure of the model is summarized in Figure 1.

**Figure 1.**
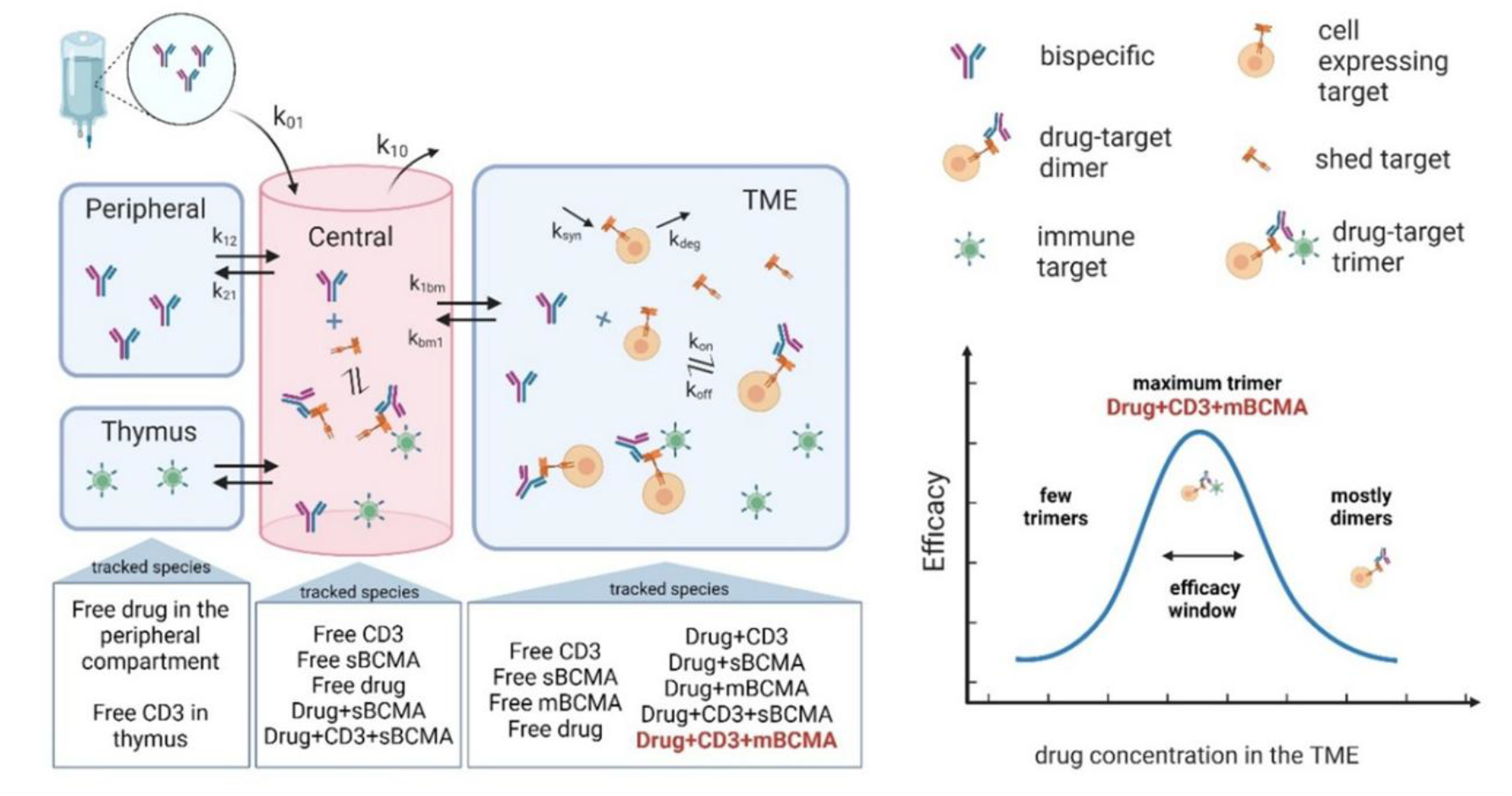
A schematic representation of the key species tracked in the model described in Supplementary Materials, and the overall objective of maximizing the pharmacologically active trimer (PAT = Drug + CD3 + mBCMA).

Teclistamab has a molecular weight of 146 kDa, with a reported dissociation constant (KD) of 28.03 nM for CD3 and 0.18 nM for BCMA (13). Its PK is well-described by a two-compartment model with linear clearance; PK parameters have been reported in (14) based on the results of the MajesTEC-1 trial in patients with relapsed/refractory MM. Additionally, PK curves and relative concentrations of the drug in the plasma and the bone marrow were reported in Girgis et al. (15), allowing estimation of parameters of drug distribution between the plasma and bone marrow compartments. Baseline concentration and turnover rates for CD3 have been reported in (10) and (11). Turnover rates of BCMA have been estimated based on reported baseline concentrations in healthy volunteers and MM patients (16), estimating serum BCMA levels in MM patients to be 175.6±242.6 ng/mL, while in healthy volunteers they are closer to 9.28±1.9 ng/mL; the authors also reported that these values were similar between newly diagnosed patients (147.6±190.8ng/mL) and those with relapsed disease (184.3±259.1 ng/mL). Given the molecular weight of BCMA of 6 kDa (17), a median soluble BCMA level of 20 nM would translate to 20 nM*6kDa = 120 ng/ml.

Furthermore, in (18), the authors report the distribution of soluble BCMA levels, which is well approximated by lognormal distribution. Molecular weight of membrane-bound BCMA is approximately 20 kDa (19). The shedding rate of BCMA has not been reported directly; however, estimates can range from 0.0023 to almost 0.4/h (20). BCMA half-life has been estimated to be approximately 24-36 h (21); internalization rates of BCMA have been estimated in (22). All nominal parameter values, along with the baseline values used for any nonzero initial conditions are summarized in Table S1.

The model is implemented in MATLAB and solved using the ode23s function. The model is first solved for a period of 50 days in the absence of treatment - the time duration was chosen arbitrarily to ensure that the pre-treatment system is fully pre-equilibrated. This pre-equilibration step allows for a determination of biomarkers that could be measured at the start of treatment. Treatment is implemented after this pre-equilibration phase. A parameter sensitivity analysis is conducted across a range of doses and schedules by assessing the following: what is the minimal fractional change in each parameter or initial condition from its nominal value that results in the area under the curve (AUC) of the PAT changing by 20%. The AUC of the PAT (in nM·d) was calculated using the trapezoidal rule over the full treatment period (including all administered doses), rather than restricting to a single steady-state dosing interval, to better capture the cumulative pharmacological exposure associated with trimer dynamics. As simulations were performed over the duration required to administer all doses, plus one additional dosing interval after the final dose, we normalize the AUC by this simulation time to have a standardized comparison across treatment protocols. All non-PK parameters were analyzed for their sensitivity, as were all nonzero initial conditions. If the AUC of the PAT does not change by 20% when a parameter is varied by 100% of its nominal value, the parameter is classified as insensitive.

The results of this sensitivity analysis informed our design of a virtual patient population. In particular, we identify a subset of the sensitive parameters and initial conditions that we assess are likely to vary across patients. Each of these attributes are assumed to follow a lognormal distribution. A virtual patient is defined by independently sampling a value of each attribute from its assumed distribution. Using this virtual population, we can assess the average teclistamab efficacy across the population, and we can also assess efficacy in subpopulations stratified by clinically measurable biomarkers. All code is available on GitHub at https://github.com/jgevertz/teclistamab.

## Results

### Baseline model behavior

First, we set out to confirm that the proposed model reproduces available average human PK data and achieves the expected steady state concentrations. In (15), the authors report teclistamab PK for 0.72 mg/kg and 1.5 mg/kg subcutaneous (SC) dose administration, and PK for 0.27 mg/kg and 0.72 mg/kg administered intravenously (iv). They also simulated PK profiles for multiple doses to assess the projected trough concentration at steady state (Ctrough, also referred to as Cmin). These simulations were benchmarked against published results from an ex vivo T-cell dependent cytotoxicity assay for teclistamab (15). In that assay, cytotoxicity was induced at EC50 of 2.53 nM (with a range of 1.47-4.21) and mean EC90 of 15.56 nM (range 1.71-41.29), where ECN refers to the concentration necessary to achieve N% of the maximum efficacy. Given teclistamab’s molecular weight of 146 kDa, this translates into average concentrations of EC50 = 369 ng/ml, EC90 = 2272 ng/ml, and max EC90 = 6028 ng/ml. Girgis et al. (15) then show that a dose of 0.72 mg/kg SC is predicted to achieve a Cmin steady state level of at least 6000 ng/ml, corresponding to reported max EC90 concentration.

In Figure 2A, we demonstrate that, at the nominal parameterization (Table S1), our model reproduces the PK profiles reported in (15) for doses of 0.72 and 1.5 mg/kg administered subcutaneously (SC). We note that while both the model proposed herein and the model in (15) both well-describe the PK data, our fit-for-purpose model additionally describes drug-target interactions in the plasma and the bone marrow, allowing us to expand the analysis beyond PK. As we expect the pharmacologically active trimer (PAT) in the bone marrow to drive efficacy, we also report the calculated value of the PAT at the nominal parameterization for a dose of 0.72 mg/kg SC and 1.5 mg/kg SC (Figure 2B).

**Figure 2.**
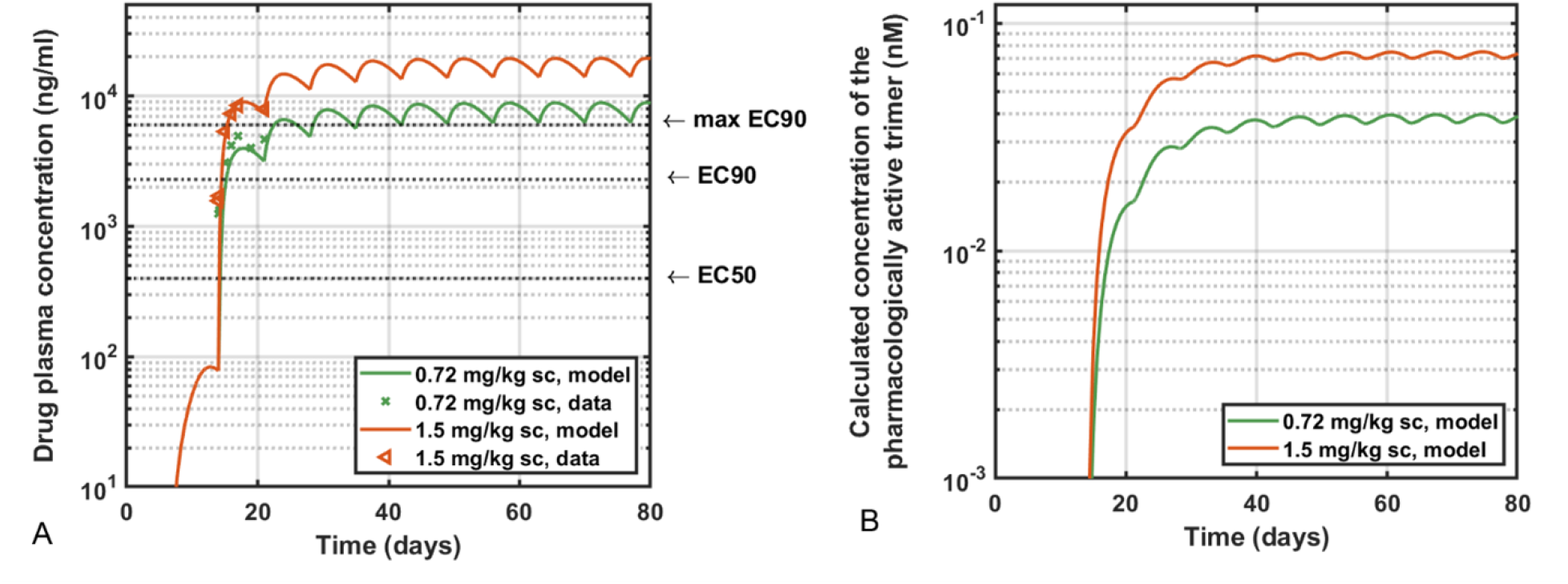
Model validation against reported PK profiles of teclistamab. (A) Simulated PK curves for administration of 0.06 mg/kg and 0.3 mg/kg loading doses, followed by full doses of either 0.72 or 1.5 mg/kg of the drug administered subcutaneously every week; PK data were digitized from (15). (B) Calculated concentrations of the pharmacologically active trimer (PAT) in the bone marrow.

### Sensitivity Analysis

Given that for a drug like teclistamab, the pharmacologically active trimer is expected to drive efficacy, we next seek to determine the model parameters that have the largest impact on PAT formation, as detailed in the Methods. To assess the impact of model parameters on drug efficacy, we chose the normalized AUC of the pharmacologically active trimer (PAT) as the key metric, since standard pharmacokinetic metrics such as Cmax or Cmin may not reliably correlate with pharmacological activity. A higher dose may lead to a higher Cmax but can also drive a faster decline in PAT levels over time due to stoichiometric imbalances. Conversely, a lower dose may achieve a lower initial Cmax but sustain higher PAT levels at later time points. Moreover, due to interpatient variability, these effects may occur at different doses and at different time points across individuals. Using the AUC of the PAT as a metric integrates these dynamic effects over time and thus provides a more robust measure of pharmacologically relevant exposure.

Figure 3 summarizes the results of this analysis, showing by how much each sensitive parameter or nonzero initial condition must be perturbed to change the normalized area under the curve (AUC) of the PAT by 20% across a range of doses and schedules. Parameters that can vary by 100% (i.e, from 0 to 2x from the nominal value) without causing the normalized AUC of the PAT to change by 20% are classified as insensitive and not included in Figure 3.

**Figure 3.**
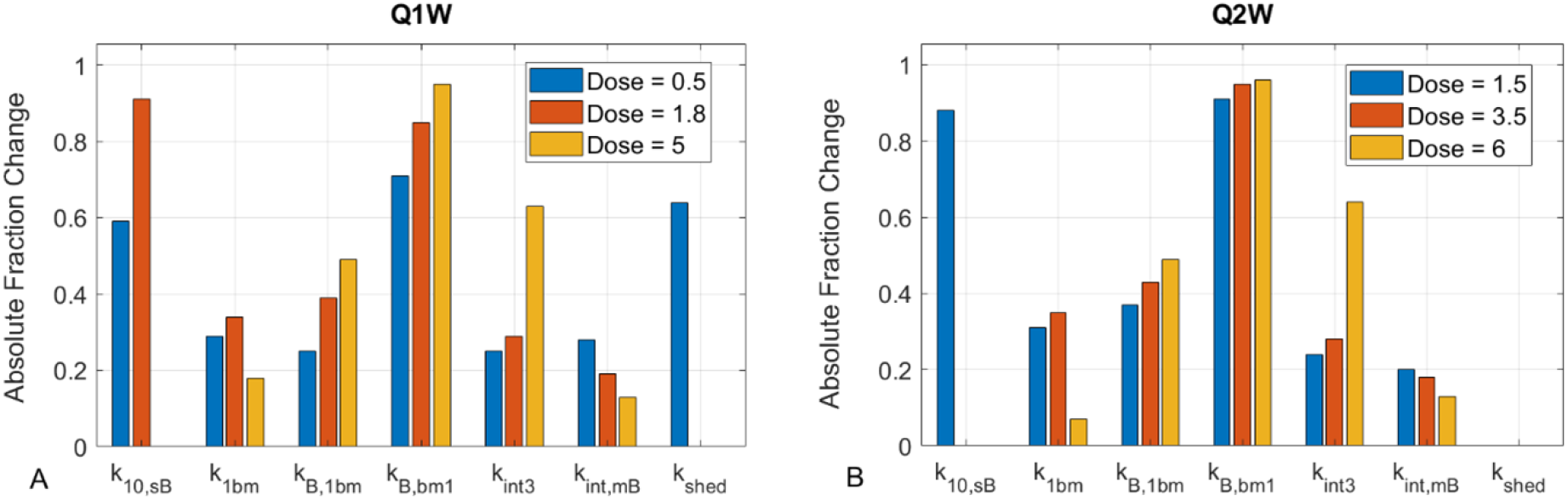
Results of the sensitivity analysis, showing by how much a parameter or a non-zero initial condition needs to be perturbed to change the AUC of the pharmacologically active trimer (PAT) by 20%. (A) Parameter sensitivity to perturbation weekly drug administration; (B) Parameter sensitivity to perturbation for Q2W administration. For each administration schedule, a dose below (blue), at (red), and above (yellow), the peak of the bell-shaped efficacy curve is chosen. If a parameter can be varied by 100% without meeting the threshold change in the AUC of the PAT, the parameter is classified as insensitive and not shown. If no bar is shown for a specific dose, that indicates that, at that dose, the parameter is classified as insensitive.

The results in Figure 3 show that, out of all the non-PK model parameters, the most sensitive ones are the internalization rates of BCMA (*k*_*int,mB*_) and CD3 (*k*_*int*3_), the rate of drug distribution into the tumor from plasma (*k*_1*bm*_), the rate of soluble BCMA distribution from plasma to bone marrow (*k*_*B*1*bm*_) and the reverse direction (*k*_*bBm*1_). For low doses only, the rate of systemic clearance from soluble BCMA (*k*_10*sB*_) and the rate of BCMA shedding (*k*_*shed*_) are also sensitive parameters. Because of the pre-equilibration, the model was insensitive to changing the value of any nonzero initial conditions.

### Virtual Population

The approach of selecting a dose based on achieving steady state of some anticipated minimally effective concentration, such as EC90, is meaningful if the drug exhibits an Emax efficacy curve, where higher concentrations do not lead to a loss of efficacy. However, for drugs like teclistamab that have a bell-shaped efficacy curve, such an approach will not be appropriate, since excessively high concentrations can result in both higher likelihood of toxicity, and in loss of efficacy. Matters are further complicated by the expectation that different individuals will maximize PAT at different “sweet spot” drug concentrations. Furthermore, because high concentrations of the drug in the TME can reduce PAT formation, it is possible that some cases of treatment relapse may stem not from acquired resistance, but from stoichiometric imbalances induced by overdosing. This possibility further motivates our exploration of biomarker-based patient stratification to optimize exposure to the pharmacologically active trimer.

To mitigate this, we next analyze a virtual population in which each patient in the population is assigned a personalized value for a subset of the most sensitive model parameters and initial conditions. Of the most sensitive parameters identified in Figure 3, the rates of target internalization (*k*_*int*3_, *k*_*int,mB*_) and the rate of soluble BCMA distribution between the plasma and bone marrow (*k*_*B*,1*bm*_ and *k*_*B,bm*1_) are the least likely to vary between patients. On the other hand, drug distribution into the tumor (*k*_1*bm*_) and target shedding (*k*_*shed*_) could vary significantly across patients. Finally, baseline expression levels of CD3 and BCMA will affect the magnitude of the formation of the pharmacologically active trimers and are expected to vary between patients.

Based on these considerations, we define a virtual patient by assigning a personalized value to each of the following characteristics: 1) baseline level of the membrane-bound BCMA in the TME, *mB*_*bm*_(0); 2) rate of BCMA shedding, *k*_*shed*_; 3) rate of drug distribution into the bone marrow, *k*_1*bm*_; and 4) baseline level of free CD3 in the TME, *CD*3_*bm*_(0). We also include variability in the drug clearance rate *Cl*_1_ as this pharmacokinetic parameter can significantly affect the availability of drug to form a pharmacologically active trimer. We assume that each of these parameters is sampled from a lognormal distribution with the mean set to the nominal value reported in Table S1 and hypothesized realistic values for the standard deviation (Figure 4). Standard deviations for most parameters were selected phenomenologically to reflect variability on the same order of magnitude as their mean values. For sBCMA-related parameters, empirical data were available from (16,18) and used to calibrate the variability to match observed distributions in patient populations. A virtual patient is then defined by randomly, and independently, sampling a parameter value from each of these five distributions.

**Figure 4.**
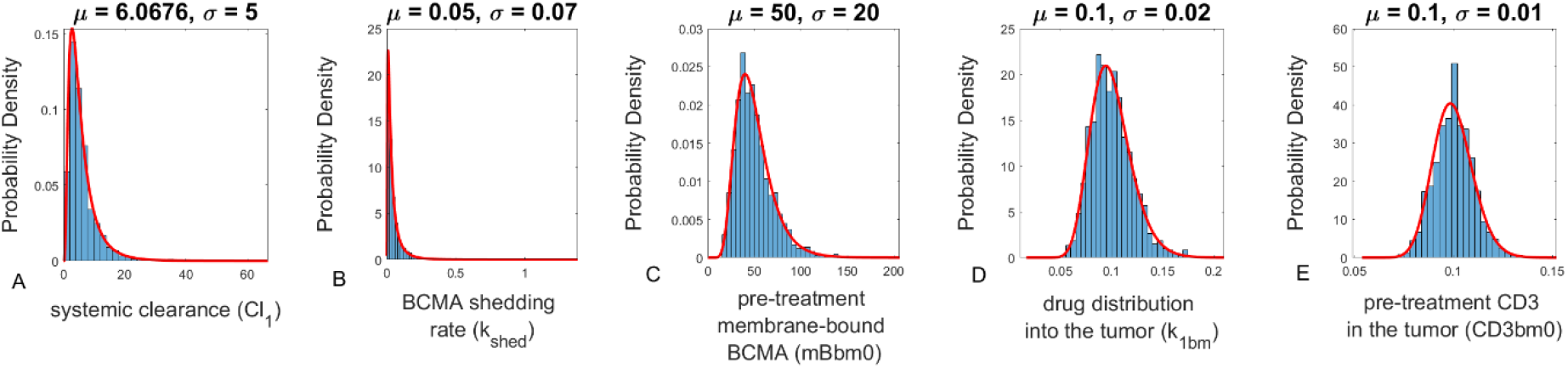
Distributions for the parameters and initial conditions that characterize a virtual patient. (A) systemic clearance, (B) BCMA shedding rate, (C) baseline expression of membrane-bound BCMA in the bone marrow, (D) intercompartmental rate constant (plasma to tumor), and (E) baseline CD3 in the TME. The blue bars represent 1200 samplings of the parameter from its theoretical distribution. Parameter ***μ*** denotes the mean value; ***σ*** denotes the standard deviation; red line denotes the theoretical distribution from which the 1200 values were sampled.

Prior to simulating treatment, we perform a pre-equilibration step for each virtual patient to reveal the distribution of two clinically measurable biomarkers: soluble BCMA in plasma and membrane-bound BCMA in the bone marrow, as well as other pre-treatment characteristics, such as baseline levels of CD3 in plasma and bone marrow, and sBCMA levels in the bone marrow (Figure 5). The predicted distribution of soluble BCMA in plasma corresponds to values that have been experimentally observed in (16), with a lognormal distribution of values among patients, as shown in (18). These measurable characteristics will be used to stratify the patients in the virtual population.

**Figure 5.**
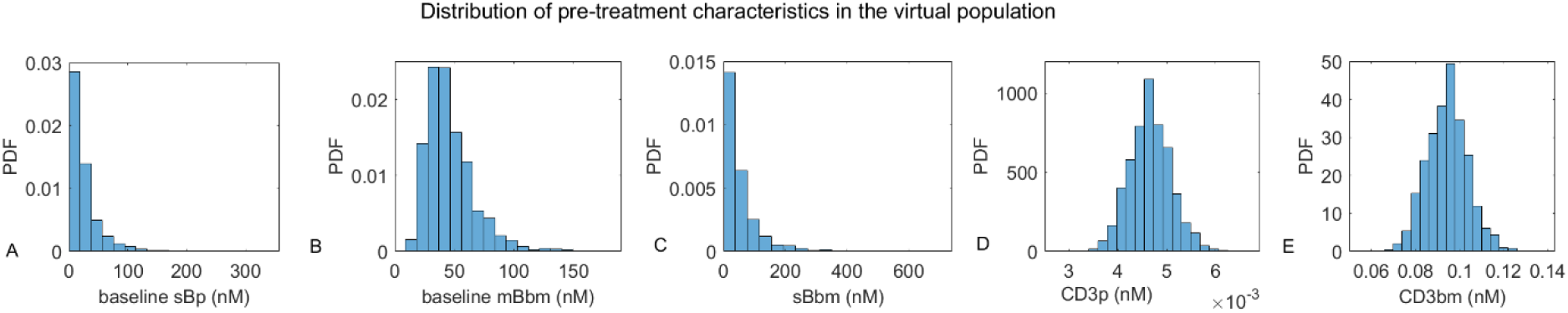
Distribution of pre-treatment characteristics in the virtual population after model-pre-equilibration. (A) baseline soluble BCMA in the plasma, (B) baseline membrane-bound BCMA in the bone marrow, (C) baseline soluble BCMA in the bone marrow, (D) baseline CD3 in the plasma, (E) baseline CD3 in the bone marrow.

In Figure 6, we show the dynamics of both free drug in the plasma and the pharmacologically active trimer across the virtual population. To determine the 95% confidence interval for each variable, the maximum and minimum value of the trajectory for each virtual patient is calculated, starting from the third administration of the drug. Any trajectory for which the maximum is in the top 2.5%, or the minimum is the bottom 2.5%, is considered outside of the 95% confidence interval. Since the virtual population is composed of 1200 virtual patients, the 95% confidence interval is described by the trajectories of the 1140 virtual patients which were not filtered out due to being extremal.

**Figure 6.**
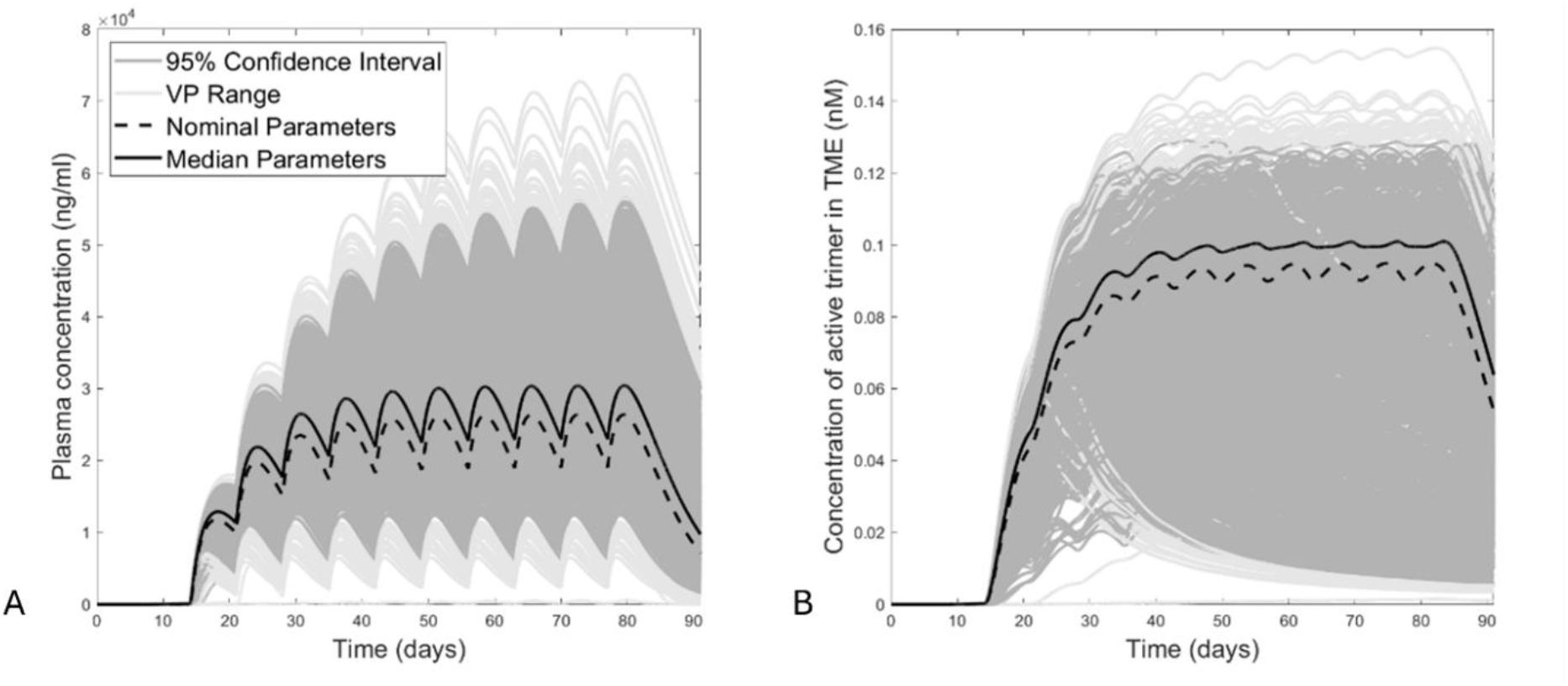
Individual trajectories across the entire virtual population (1200 virtual patients) at a dose of 2 mg/kg. (A) Plasma drug concentration. (B) Pharmacologically active trimer (PAT) The diversity of response profiles highlights patient-to-patient variability in PAT dynamics.

### Analysis of Dose Response in Full Population

To find the dose that will maximize the normalized AUC of the PAT across the virtual population, we conduct a dose sweep by varying doses from 0.1 to 10 mg/kg, administered either weekly (Q1W) or every two weeks (Q2W). This range specifically includes doses that were previously tested for teclistamab, including 0.27 mg/kg; 0.72 mg/kg; 1.5 mg/kg and 2.16 mg/kg (15). We also include other values between and up to an upper threshold of 10 mg/kg to computationally evaluate larger doses than could be feasibly tested in the clinic (doses up to 6 mg/kg were assessed in (13)). For each dose and schedule, we compute the median value of the normalized AUC of the pharmacologically active trimer across a virtual population of 1200 patients. This number was chosen to allow for sufficient variability and to mimic the possible size of a Phase 3 trial. Notably, because the absolute threshold of normalized pharmacologically active trimer required for efficacy is unknown, dose optimization focuses on maximizing relative PAT AUC rather than achieving a predefined target value.

Figure 7A shows that there exists a clear “sweet spot” in the dosing space, with the median normalized AUC of the PAT (and thus the predicted efficacy) achieving its absolute maximum at an intermediate drug concentration. The figure also identifies what we term the “near-optimal dose range”, defined as drug concentrations that give a median normalized AUC within 10% of the absolute maximum AUC value. We find that the optimal average dose for Q1W administration is 1.8 mg/kg, with a near optimal range of 1.3 and 2.5 mg/kg. For the Q2W protocol, the optimal dose identified is 3 mg/kg, with a near optimal range of 2.5 to 4 mg/kg. The true recommended phase II dose (RP2D) for teclistamab is 1.5 mg/kg SC Q1W (14), preceded by step-up doses of 0.06 and 0.3 mg/kg. Thus, the proposed semi-mechanistic model can reach a comparable conclusion about the RP2D using an analysis of the AUC of the pharmacologically active trimer in a virtual population.

**Figure 7.**
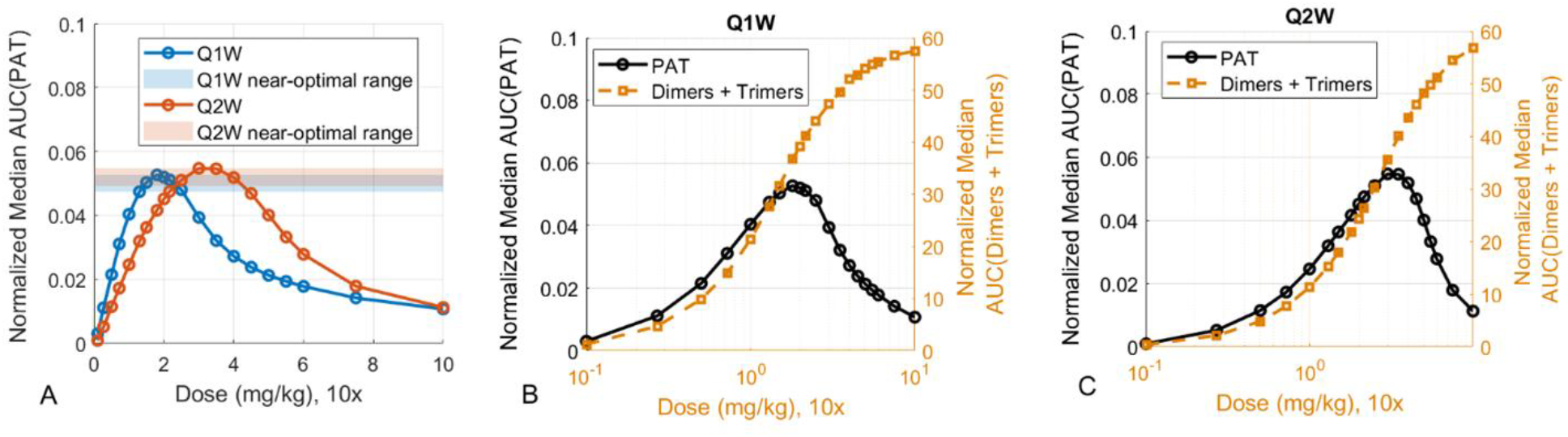
Population-level analysis of doses expected to maximize the normalized AUC of the PAT. (A) Median normalized AUC (across 1200 virtual patients) of the calculated PAT for a range of doses from 0.1 to 10 mg/kg, administered subcutaneously either Q1W (blue) or Q2W (red); shaded regions represent near-optimal dosing range, defined as being within 10% of the maximum value of the normalized AUC of the PAT. (B) PATs (left axis) and the sum of dimers and trimers (right axis) for Q1W and (C) Q2W dosing. Note that the black PAT curve in (B) corresponds to the blue curve in (A) and that the orange PAT curve in (C) corresponds to the red PAT curve in (A). Note the log x-axis. Normalized AUC has units nM.

In Figure 7B and C we plot the population-level normalized AUC of PATs (left y-axis) along with the normalized AUC of the sum of both dimers and trimers (right y-axis), for Q1W and Q2W dose administrations, respectively. These plots reveal that if both dimers and trimers contributed to pharmacological efficacy, we would have observed the standard Emax efficacy curve (red, Figure 7B and C), with efficacy increasing monotonically as a function of drug concentration. Consequently, a standard Cmin-based dose selection strategy would have been justified. However, if only the PAT drives efficacy, then we observe a clear bell-shaped curve (blue, Figure 7B and C), and thus an alternative dose selection strategy is warranted.

### Two Group Subpopulation Analysis

Next, we investigate whether we can identify subgroups of patients that may be more or less likely to respond to the recommended population-level dose of teclistamab. We start by stratifying virtual patients into subpopulations based on the clinically measurable level of soluble BCMA in the plasma (*sB*_*p*_) at the start of the treatment, as calculated at the end of the pre-equilibration step described in the Methods. Specifically, we subdivide virtual patients based on whether their level was above or below the median level of soluble BCMA (Figure 8A).

**Figure 8.**
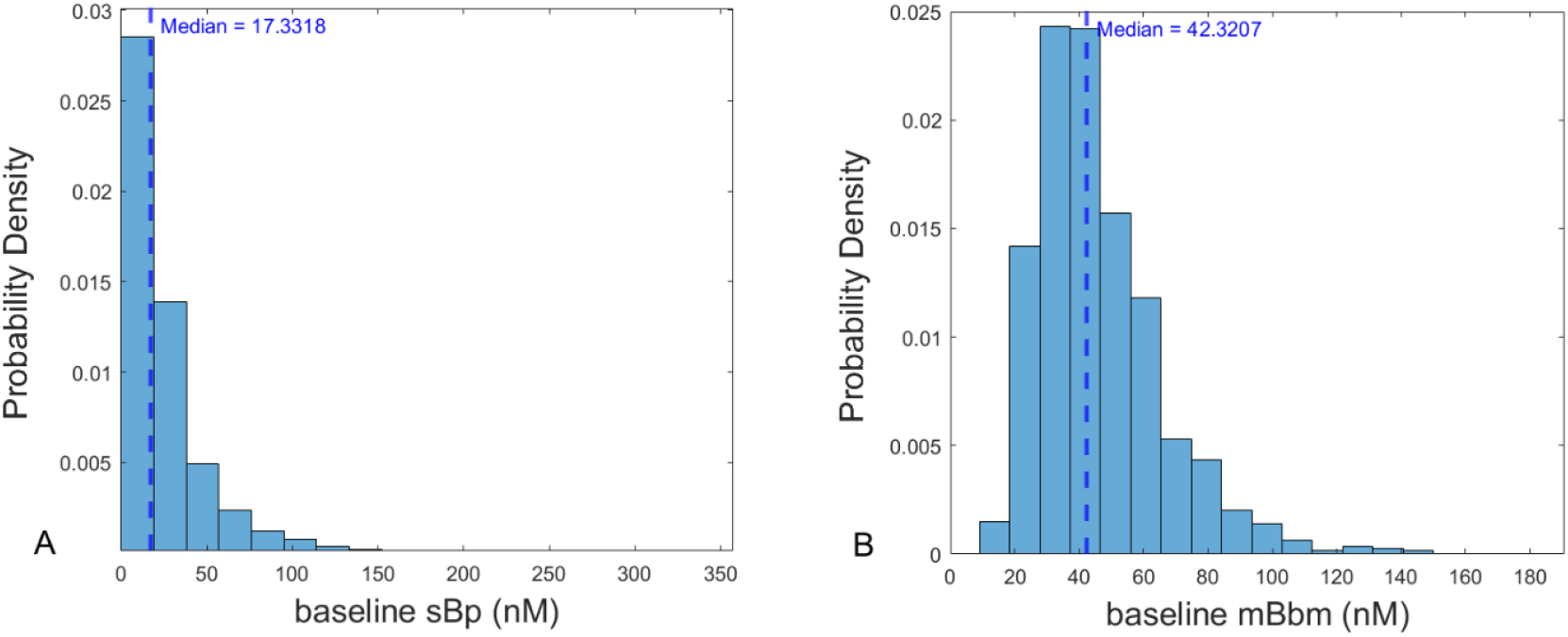
Criteria used for patient stratification. (A) Pre-treatment distribution in the virtual population of soluble BCMA in the plasma, and (B) membrane-bound BCMA in the bone marrow. The median value for each characteristic is given by blue dashed line.

For the subpopulation with levels of soluble BCMA below the full population median (low *sBp*), the median AUC of the PAT is maximized between doses of 1.3 to 2.16 mg/kg, with the peak occurring around 1.5 mg/kg, for Q1W dosing. For Q2W, the optimal dosing range is between 2.16 and 3.5 mg/kg, with the peak occurring at 3 mg/kg (Figure 9).

**Figure 9.**
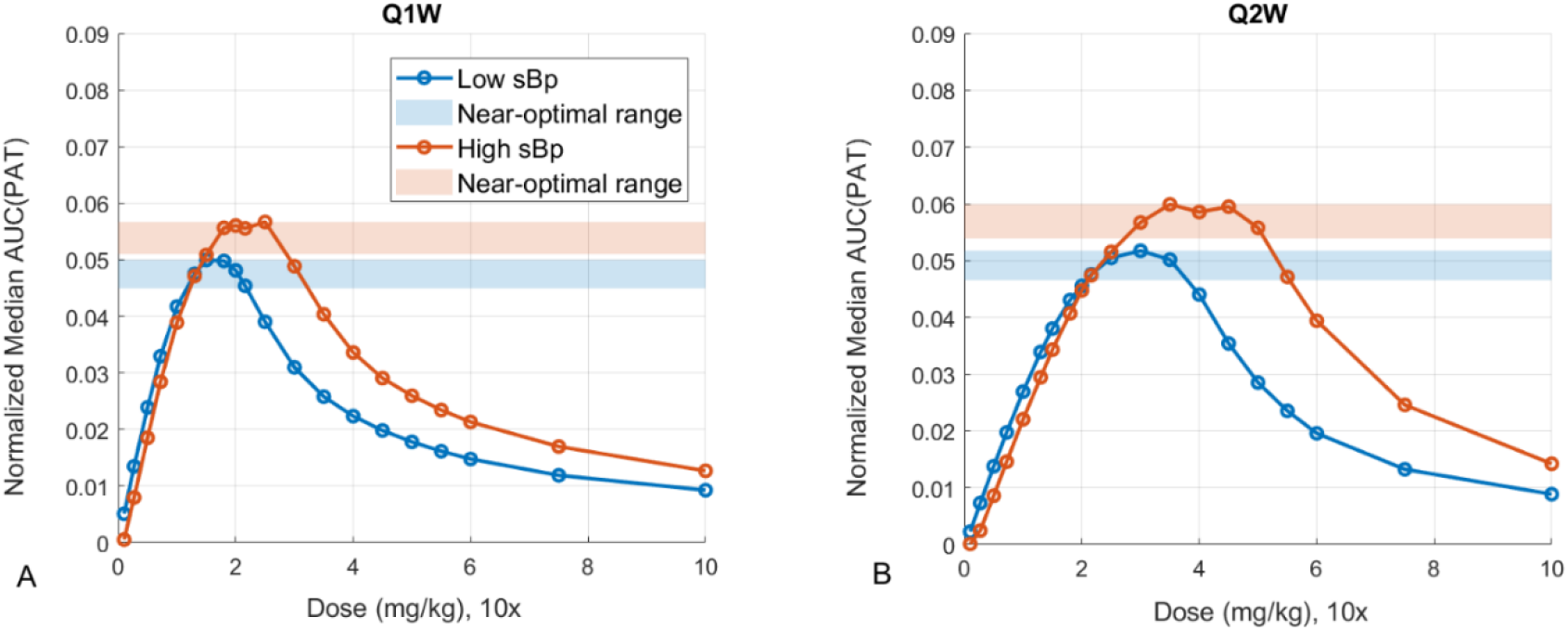
Patient stratification based on baseline levels of soluble BCMA in the plasma (***sB***_***p***_); high vs low characteristics are assigned based on whether the individual’s values fall above or below the median in the population. Median normalized AUC for a subpopulation is shown for (A) Q1W schedule; (B) Q2W schedule. Shaded regions represent near-optimal dosing range, defined as being within 10% of the maximum value of the normalized AUC of the PAT. Normalized AUC has units nM.

For the high *sBp* subpopulation, the median AUC of the PAT is maximized between doses of 1.5-2.5 mg/kg, with the peak at 2.5 mg/kg, for Q1W. For Q2W, the optimal dosing range is between 3 and 5 mg/kg with the peak at 3.5 mg/kg. Thus, for those VPs with higher baseline soluble BCMA levels in the plasma (high *sBp*), treatment efficacy could be increased at higher doses than selected for the average. It should be noted that this analysis focused solely on maximizing expected efficacy and does not consider possible toxicity that could result from these higher doses. The summary of dose predictions for the full population, and subdividing into subpopulations based on sBCMA, is given in Table S2. The impact of this approach to patient stratification on individual trajectories is shown in Figure 10, compared to the full population (Figure 7A).

**Figure 10.**
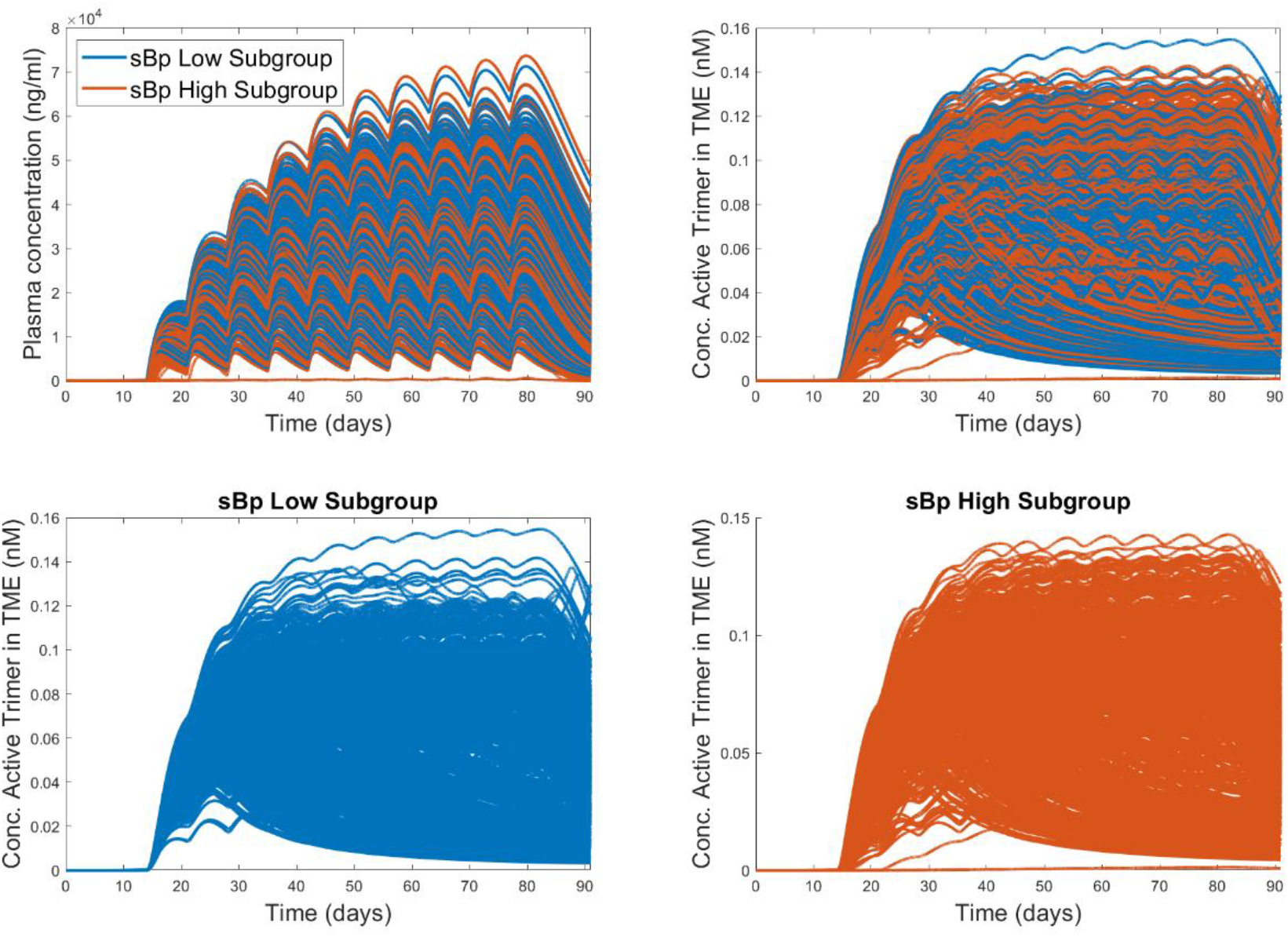
PAT trajectories for the full virtual population stratified into two subgroups based on soluble BCMA levels, ***sBp***, at baseline.

### Four Group Subpopulation Analysis

Next, we assess whether any additional insights can be obtained if the patient population is further stratified based on both baseline levels of soluble BCMA in the plasma (Figure 8A) and the baseline expression level of membrane-bound BCMA in the bone marrow (Figure 8B) which conceivably could be assessed during pre-treatment biopsy.

This further patient stratification leads to several interesting results. First, for the Q1W dosing, we see that not all subpopulations maximize the pharmacologically active trimer at the same doses, with those with low membrane-bound BCMA in the TME requiring lower doses in the range of 1-2 mg/kg (Figure 11A, C), and those with high membrane-bound BCMA (Figure 11B, D) maximizing the normalized AUC of the PAT at doses in the range of 1.5-3 mg/kg.

**Figure 11.**
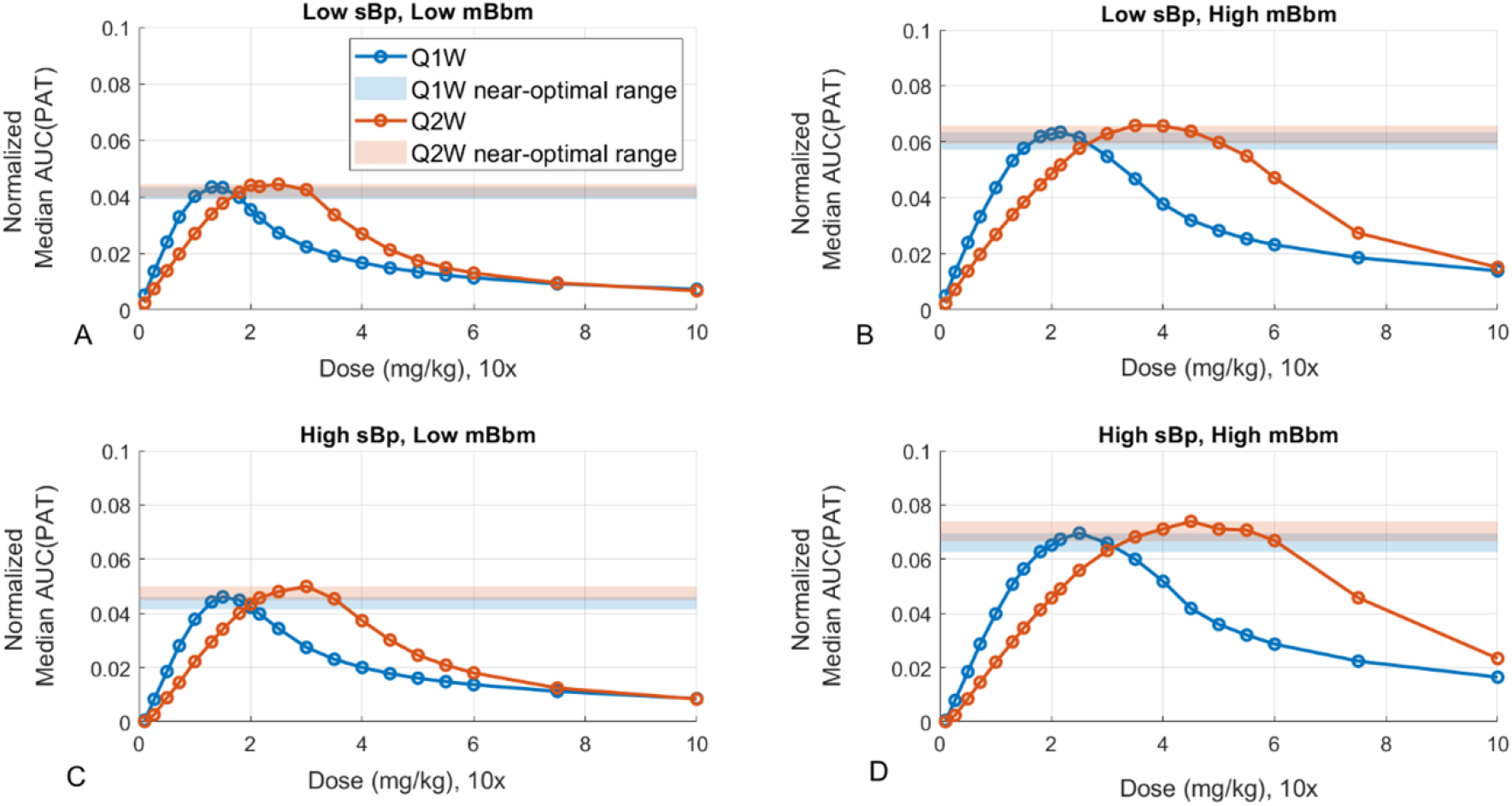
Patient stratification based on baseline levels of both soluble BCMA in the plasma (***sB***_***p***_), and membrane-bound BCMA in the TME (***mB***_***bm***_); high vs low characteristics are assigned based on whether the individual’s values fall above or below the median in the population. (A) Median normalized AUC for a subpopulation with low sBCMA in plasma and low mBCMA in the bone marrow. (B) Median normalized AUC for a subpopulation with low sBCMA in plasma and high mBCMA in the bone marrow. (C) Median normalized AUC for a subpopulation with high sBCMA in plasma and low mBCMA in the bone marrow. (D) Median normalized AUC for a subpopulation with high sBCMA in plasma and high mBCMA in the bone marrow. Shaded regions represent near-optimal dosing range, defined as being within 10% of the maximum value of the normalized AUC of the PAT. Normalized AUC has units nM.

Additionally, if in the beginning of the trial one was unclear as to which patients would be most likely to respond, and if the goal were to find those patients for whom the value of the trimer would be maximized, this approach would allow for the identification of such “no regrets” patients. This is similar to a “no regrets” dosing approach, where if a dose does not lead to efficacy despite fully engaging the mechanism, then targeting the selected mechanism might not be appropriate for a given indication. Here, those who are most likely to respond (assuming direct correlation between high PAT and efficacy) are patients with high soluble and membrane-bound BCMA, followed by high membrane-bound and low soluble BCMA, followed by those with high soluble BCMA and low membrane-bound BCMA, and finally those with low soluble and membrane-bound BCMA. Interestingly, there appears to be a minimal difference between these subgroups up to 1 mg/kg Q1W and up to 1.8 mg/kg Q2W (Figure 12).

**Figure 12.**
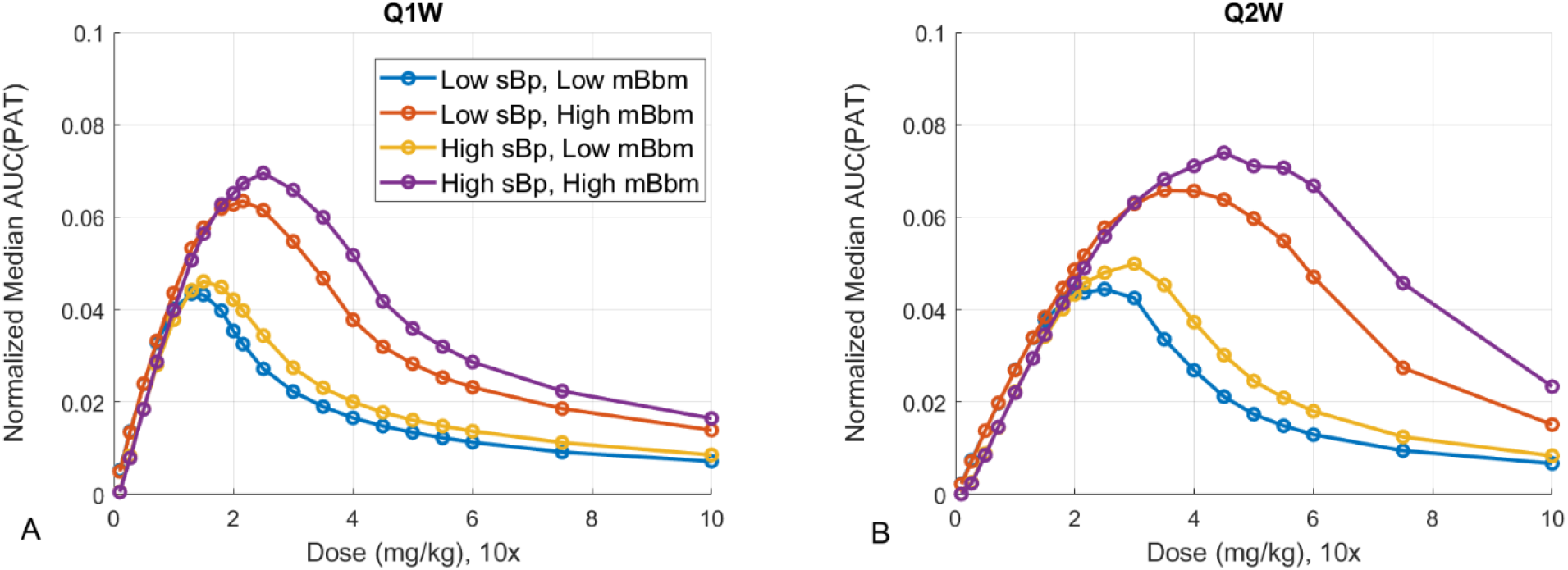
Patient stratification based on baseline levels of both soluble BCMA in the plasma (***sB***_***p***_), and membrane-bound BCMA in the TME (***mB***_***bm***_); high vs low characteristics are assigned based on whether the individual’s values fall above or below the median in the population. (A) Q1W dosing regimen. (B) Q2W dosing regimen Normalized AUC has units nM.

Overall, the Q1W schedule is predicted to have a lower variability of response as all the median subpopulation AUCs are closer together at the peak, compared to much larger variability between subpopulations for Q2W dosing. The summary of dose predictions for this stratification of patients is given in Table S2, and individual PAT trajectories are shown in Figure 13.

**Figure 13.**
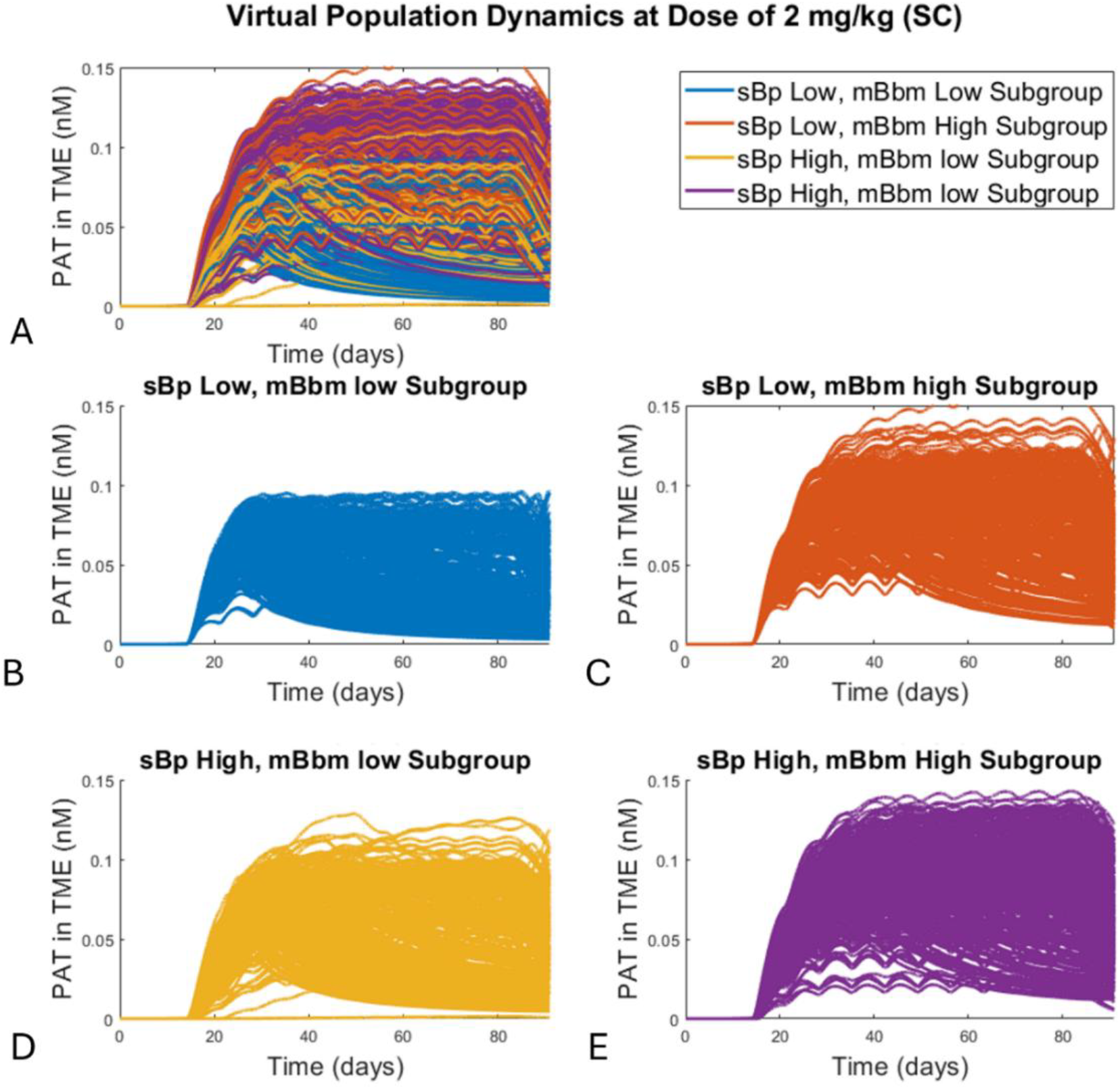
PAT trajectories for the full virtual population further stratified into four subgroups based on both soluble BCMA and membrane-bound BCMA levels at baseline. (A) PAT in the bone marrow for the entire population. (B) PAT trajectories for the subgroup with low pre-treatment levels of soluble BCMA in plasma and low membrane-bound BCMA in the bone marrow (low/low). (C) PAT trajectories for the low/high subgroup. (D) PAT trajectories for the high/low subgroup. (E) PAT trajectories for the high/high subgroup. High and low classifications are determined by whether pre-treatment levels of sBCMA in plasma or mBCMA in the bone marrow were above or below the population median.

## Discussion

Dose optimization is a challenging and time-consuming process, and can be as much of an art as it is science. It can become even more complicated for bispecific antibodies that have a bell-shaped, rather than an Emax, efficacy curve (10,11). These include most bispecific T cell engagers, whose mechanism of action is predicated on the drug binding to both targets simultaneously, forming a pharmacologically active trimer. The particular challenge of dose optimization for these compounds lies in the fact that, for low drug concentrations, few pharmacologically active trimers are formed. However, if the doses are too high, the stoichiometry shifts away from trimers and towards increased dimer formation, diminishing efficacy. Taken together, this results in the aforementioned “bell-shape” curve, with the lose-lose situation that over-dosing can lead to both excess toxicity and a loss of efficacy.

Optimal dosing of such bispecifics is further complicated by patient heterogeneity. Individual patient characteristics, which may affect drug absorption, accumulation, distribution, as well as different levels of target expression, will necessarily impact how much of the pharmacologically active trimer can be formed. As a result, a dose that is optimal for a patient with one combination of these characteristics may be suboptimal for a patient with different features. Furthermore, administering a dose that is too high for a particular patient may appear as emergence of treatment resistance; however, in his case it may be a result of over-dosing, where drug-target stoichiometry is shifted away from maximizing PAT formation, rather than the typically irreversible genetic resistance. Fortunately, in this case, sensitivity to the drug may be restored by reducing the dose. Stratified dosing approaches based on measurable biomarkers could help mitigate this risk and further improve patient outcomes.

Here we explore a computational approach to the bispecific dose optimization problem. We use as a test example a model of teclistamab, a bispecific antibody used for treatment of multiple myeloma that targets CD3 on the immune cells on one arm, and BCMA on cancer cells on the other arm. For teclistamab, the PAT is formed by the drug bound to both CD3 and membrane-bound form of BCMA in the bone marrow. We consider a virtual population, where patient variability is captured by determining the most sensitive model parameters, and assigning each virtual patient a value of these parameters randomly sampled from a lognormal distribution. We first identify the dose and schedule (either Q1W or Q2W) for which the pharmacologically active trimer in the TME is maximized for the median patient in the full population. Using the normalized AUC of the PAT as our metric, we were able to rediscover that the recommended phase II doses for teclistamab fall within the predicted optimal range. Specifically, we found that the median AUC of the PAT was maximized for doses between 1.3 and 2.5 mg/kg when administered weekly (Q1W) and between 2.5 and 4 mg/kg when administered biweekly (Q2W). The recommended Phase 2 dose (RP2D) is a weekly subcutaneous dose of 1.5 mg/kg (14), with follow up studies suggesting that switching to 1.5 mg/kg on a Q2W schedule can maintain durable responses for individuals who responded to the original dosing protocol (24).

Next, we assess whether dose projections could be refined if we subdivide patient populations based on certain characteristics that may be measurable in the clinic, particularly the baseline levels of both soluble BCMA in the plasma and membrane-bound BCMA in the TME. We found that, predictably, patients with low soluble BCMA (defined as having baseline sBCMA concentrations below the median in the virtual population) are predicted to have lower PAT AUC compared to those with high soluble BCMA (defined as above the median), although the doses that maximize PAT are estimated to be in a similar range. Based on our analysis, it is possible that the strength of both response and adverse events might be greater for those with high soluble BCMA than for those with low concentrations, since high levels of soluble BCMA, while acting as a sink, are also likely to be proxy for severity of disease and availability of the pharmacologically active target in the TME. This highlights the potential utility of monitoring sBCMA levels to improve patient stratification and guide therapeutic decisions, such as the adjustment of dosages or combining therapies to overcome resistance mechanisms.

Stratifying patients into four subgroups based on high/low levels of soluble BMCA in the plasma and high/low levels of membrane-bound BCMA in the TME showed more differences between the subgroups with respect to the “optimal” doses. Low/low subpopulations are found to receive maximum benefit at lower drug doses compared to the high/high subpopulations. However, the high/high subpopulation is predicted to have a more robust response to treatment, as measured by the values of the normalized AUC of the pharmacologically active trimer. Interestingly, the variability of response is predicted to be greater for Q2W dosing regimen than for Q1W.

This subpopulation approach provides a middle ground between the designing protocols for the average patient and personalizing therapy to the individual. Currently, a more “blanket” dosing approach is generally used, where the same dose is administered to everyone who meets the general inclusion criteria, or is adjusted mostly based on covariates, such as weight or gender. On the other end of the spectrum is personalized medicine, which remains elusive due to numerous technological, logistical, economical, and regulatory barriers (25,26), and thus is often infeasible on a large scale. Here, we show virtual population analysis can assist in identifying subpopulations that may be more likely to respond, or those who may benefit from, different dosing strategies based on their individual, measurable characteristics. Perhaps such “semi-personalized” medicine may be a more technically, logistically and economically feasible option for leveraging modeling and simulation to improve patient outcomes.

## Acknowledgements

JLG acknowledges use of the ELSA high-performance computing cluster at The College of New Jersey for conducting the research reported in this paper. This cluster is funded in part by the National Science Foundation under grant numbers OAC-1826915 and OAC-1828163.

## Author Contributions

Both authors contributed equally to writing the manuscript, designing research, performing research, and analyzing data.

## Code availability

All code is available on GitHub at https://github.com/jgevertz/teclistamab.

## Supplementary Information

### Model description

Teclistamab is a bispecific antibody that can bind two targets: CD3 expressed on T cells, and BCMA expressed particularly on multiple myeloma (MM) cells in the bone marrow (the TME). BCMA is a membrane-bound target (mBCMA) but it can shed, meaning that it can be cleaved from cells and found in its soluble form (sBCMA) in both the plasma and the TME. Shed BCMA is not expected to affect anti-tumor activity and instead is expected to act as a sink. Here we detail PK/PD model we have developed of teclistamab, the structure of which is briefly summarized in Figure 1 of the main text.

T cells are produced in the thymus, so we assume that free CD3 (*CD*3_*th*_) is produced in the thymus compartment at a rate *k*_*syn,th*_, and can distribute from thymus to the plasma compartment at a rate constant *k*_*th*1_. This process is described by the following equation:

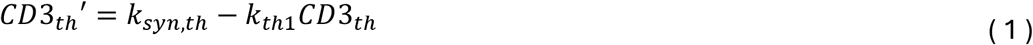

In the plasma compartment, free CD3 (*CD*3_*p*_) can be cleared through internalization at a rate *k*_*int*3_, and can reversibly bind to free drug in the plasma *D*_*p*_ to form a dimer *D*3_*p*_, with association and disassociation rates *k*_*on*3,*p*_, *k*_*off*3,*p*_, respectively. Free CD3 can also reversibly bind at the same rates to the drug-sBCMA dimer, forming the drug-CD3-sBCMA trimer (*DsB*3_*p*_). Finally, free CD3 can distribute to and from the bone marrow *CD*3_*bm*_ at rates *k*_3,1*bm*_ and *k*_3,*bm*1_, respectively. Notably, it is understood that CD3 is expressed on the T cells but for the purposes of this analysis, a more detailed description of T cell dynamics is unnecessary; therefore, here free CD3 is used as proxy for T cells expressing this receptor. The dynamics of free CD3 in the plasma compartment are described by the following equation:

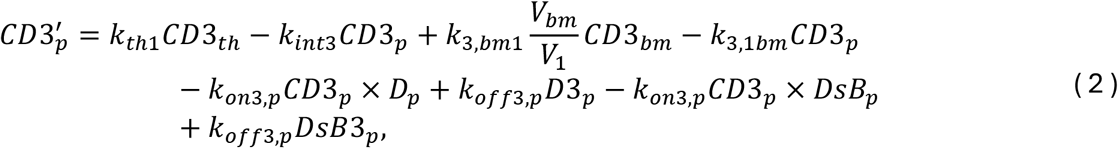

where *V*_1_ is the volume of distribution of the plasma compartment and *V*_*bm*_ is the volume of distribution of the bone marrow compartment.

Soluble BCMA in plasma, *sB*_*p*_, is assumed to primarily distribute from and back to the bone marrow (*sB*_*bm*_ at rate *k*_*B,bm*1_) and to be produced in the plasma in negligible quantities. It is assumed to clear at a rate *k*_10,*sB*_, and bind reversibly to either the free drug *D*_*p*_, or to drug-CD3 complex *D*3_*p*_. These processes are described by the following equation:

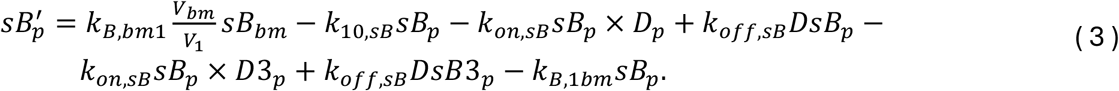

The dynamics of the free drug in plasma are described as follows. It is assumed that the drug can be administered either subcutaneously (SC) as *D*_*sc*_ and absorbed at a rate *k*_01_, or administered directly into plasma intravenously (IV). Teclistamab PK is best described by a two-compartment model (1), so we assume that drug in plasma *D*_*p*_ can distribute into the peripheral compartment *D*_*per*_ (with volume of distribution *V*_2_), or into the bone marrow as *D*_*bm*_. We assume that negligible drug-target binding occurs in the peripheral compartment. In the plasma, free drug *D*_*p*_ can reversibly bind to either free target CD3, or to soluble BCMA. It can form drug-target dimer complexes *D*3_*p*_ (drug + CD3) and *DsB*_*p*_ (drug + soluble BCMA); these complexes can in turn bind to the remaining free target to form the pharmacologically inactive trimer *DsB*3_*p*_. We assume this is inactive because the soluble form to BCMA cannot signal the tumor cell; only the membrane-bound form in the TME can. These processes are described by the following equations:

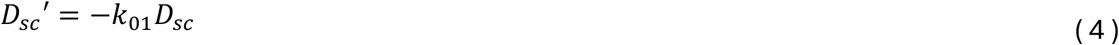

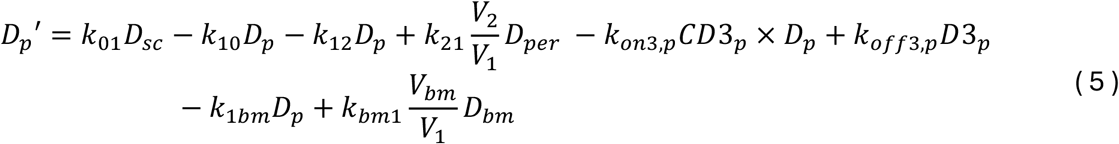

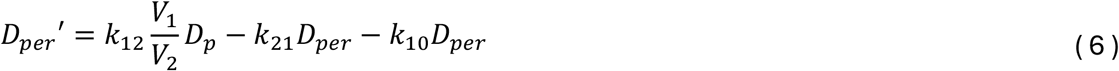

The dynamics of the drug-target complexes in plasma are described by the following equations.

Dimer: drug + soluble BCMA in plasma

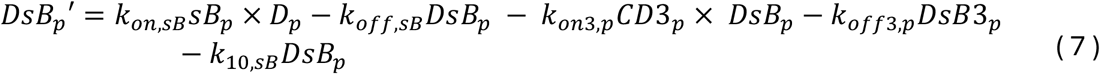

Dimer: drug + CD3 in plasma

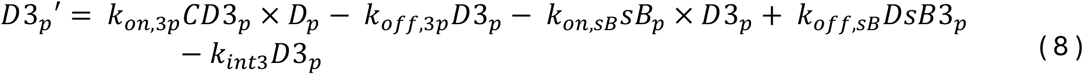

Pharmacologically inactive trimer: drug + CD3 + sBCMA in plasma

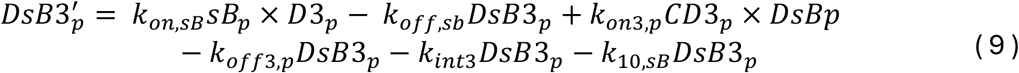

In the bone marrow, we track the dynamics of the following species: free drug (*D*_*bm*_), which is assumed to have distributed from the plasma compartment; free CD3 (*CD*3_*bm*_); free membrane-bound BCMA (*mB*_*bm*_), which can shed and produce soluble BCMA (*sB*_*bm*_). We assume that the drug–sBCMA complex remains confined to the central compartment, as its large size (∼150–160 kDa) would significantly limit transport into bone marrow tissue. This assumption is consistent with standard modeling practices for antibody–antigen complexes (2). We also track the dynamics of all the dimers (drug + CD3; drug + sBCMA; drug + mBCMA) and trimers (drug + CD3 + sBCMA and drug + CD3 + mBCMA). We assume that drug + CD3 + mBCMA is the only efficacious form of the trimer (pharmacologically active trimer, or PAT). The equations for these species are derived similarly to the ones in the plasma and are as follows.

Free membrane-bound BCMA in the bone marrow:

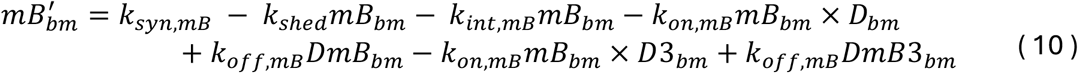

Free soluble BCMA in the bone marrow:

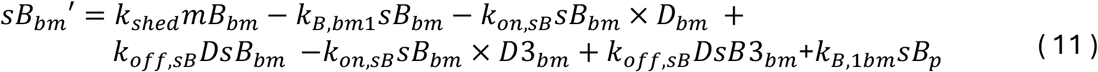

Free CD3 in the bone marrow:

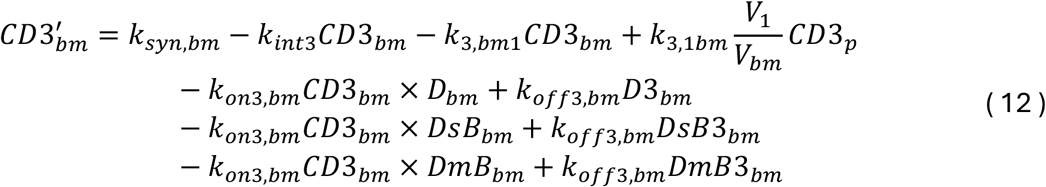

Free drug in the bone marrow:

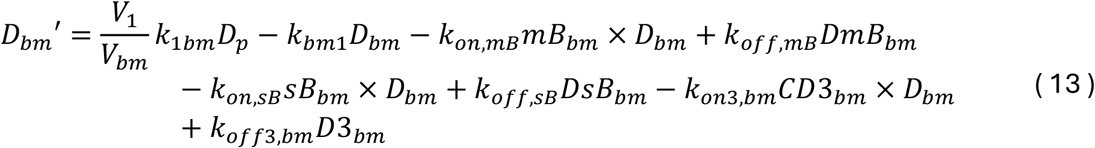

The equations for the dimer and trimer complexes in the bone marrow are as follows.

Dimer: drug + CD3 in the bone marrow:

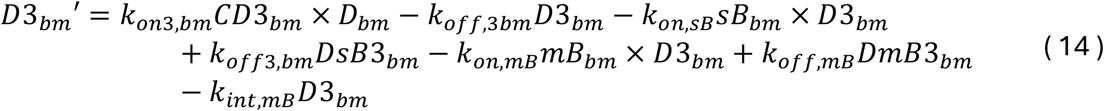

Dimer: drug+ soluble BCMA in the bone marrow:

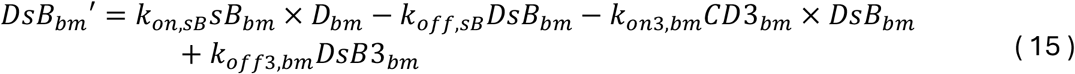

Dimer: drug + membrane-bound BCMA in the bone marrow:

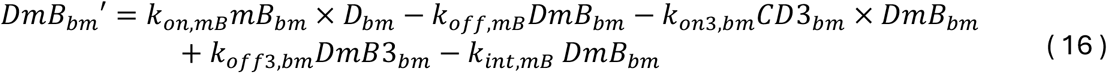

Inactive trimer: drug + soluble BCMA + CD3:

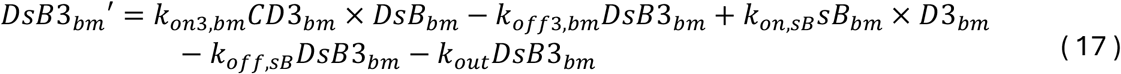

Active trimer: drug + membrane-bound BCMA + CD3:

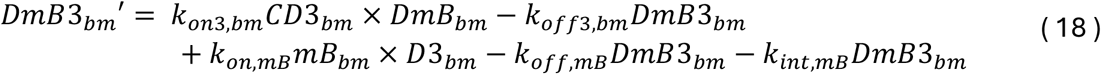

We note a few key assumptions made in constructing this model. First, since teclistamab pharmacokinetics have not demonstrated substantial dependence on mild-to-moderate renal impairment (3), renal function variability was not explicitly modeled. Second, internalization of the pharmacologically active trimer (drug + CD3 + mBCMA) is assumed to result in simultaneous clearance of the drug, membrane-bound BCMA (mBCMA), and CD3. This simplification is consistent with prior semi-mechanistic models of T-cell engagers (e.g., (4)). We acknowledge that biologically, the internalization kinetics of mBCMA and CD3 differ, with mBCMA exhibiting slower turnover relative to CD3. However, because BCMA expression was modeled with an ongoing synthesis rate, and the total available BCMA is in relative excess compared to drug levels, the effect of this assumption on pharmacodynamically active trimer formation is expected to be minimal. Future work could explore decoupling target internalization rates to better capture long-term receptor dynamics if needed.

In summary, since we assume that the pharmacologically active trimer *DmB*3_*bm*_ drives efficacy, our dose selection goal is to maximize its concentration in the TME.

## Supplementary Tables

**Table S1.** Summary of parameter values and nonzero initial conditions.

| Parameter | Description | Value | Units | Ref. |
| --- | --- | --- | --- | --- |
| **PK parameters** |  |  |  |  |
| **V<sub>1</sub>** | Volume of distribution of the central compartment | 3-4 | L | Estimated based on data reported in (1) and in (5) |
| **V<sub>2</sub>** | Volume of distribution of the peripheral compartment | 1.34 | L |  |
| **Cl<sub>1</sub>** | Clearance from the central compartment; time-dependent clearance is described as $Cl_1 = Cl_{var} e^{-\delta t}$, where $Cl_{var}$ is taken from (5) and $\delta$ is estimated to be 0.1 | 0.5 | L/d | |
| **Cl<sub>2</sub>** | Clearance from the peripheral compartment | 0.4 | L/d |  |
| **k<sub>10</sub>** | Rate constant for free drug elimination from plasma compartment; calculated as $Cl_1/V_1$ | 0.17 | 1/d | |
| **k<sub>12</sub>** | Rate constant for drug distribution into the peripheral compartment, calculated as $Cl_2/V_1$ | 0.13 | 1/d | |
| **k<sub>21</sub>** | Rate constant for drug distribution from the peripheral compartment, calculated as $Cl_2/V_2$ | 0.3 | 1/d | |
| **k<sub>10,sB</sub>** | Rate of systemic clearance of soluble BCMA | 0.04 | 1/d | Estimated to capture ranges reported in (6) |
| **KD<sub>CD3</sub>** | Dissociation constant for CD3 | 28.03 | nM | (3) |
| **KD<sub>BCMA</sub>** | Dissociation constant for CD3 for BCMA | 0.18 | nM |  |
| **k<sub>off3,p</sub>** | Dissociation rate constant for CD3; association rate is calculated as $k_{on3,p} = k_{off3,p}/KD_{CD3}$ | 285 | 1/nM/d | (7) |
| **k<sub>off,sB</sub>** | Dissociation rate constant for BCMA; association rate is calculated as $k_{onsB} = k_{offsB}/KD_{BCMA}$ | 20 | 1/d | n/a |
| **k<sub>off,mB</sub>** | Dissociation rate constant for drug-membrane BCMA complex | 20 | 1/nM/d | n/a |
| **k<sub>on,mB</sub>** | Association rate constant for drug-membrane BCMA complex | 111 | 1/d | n/a |
| $k_{1bm}$ | Intercompartmental rate constant of drug distribution from plasma into the bone marrow | 0.1 | 1/d | Calculated based on patient data from (1) |
| $k_{bm1}$ | Intercompartmental rate constant of drug distribution from bone marrow into plasma | 0.001 | 1/d | |
| $k_{B,bm1}$ | Rate of soluble BCMA distribution from BM to plasma | 0.1 | 1/d | |
| $k_{B,1bm}$ | Rate of soluble BCMA distribution from plasma to BM | 0.1 | 1/d | |
| $k_{out}$ | Clearance rate of trimer/dimer complexes from BM | 0.01 | 1/d | |
| $k_{on3,p}$ | Association rate constant for drug-CD3 complex in plasma | 10 | 1/nM/d | |
| $k_{on,sB}$ | Association rate constant for drug-soluble BCMA complex | 111 | 1/nM/d | |
| $k_{01}$ | Absorption rate constant for subcutaneous drug administration | 0.2 | 1/d | |
| $V_{bm}$ | Volume of distribution of the bone marrow compartment | 2.2 | L | |
| CD3 parameters |  |  |  |  |
| $k_{syn,th}$ | CD3 synthesis rate in the thymus; it is implicitly assumed that CD3 target is expressed on T cells, which are produced in the thymus and migrate into other compartments | 0.0005 | nM/d | Estimated based on parameters provided in (4,7) |
| $k_{th1}$ | Rate of CD3 distribution from thymus into the plasma compartment | 0.5 | 1/d | |
| $k_{int3}$ | CD3 internalization rate | 16.6 | 1/d | |
| $k_{syn,bm}$ | Rate of CD3 synthesis in the bone marrow proportional to T cell proliferation at the site of action | 19.8 | nM/d | |
| $k_{3,bm1}$ | Intercompartmental rate constant (tumor to plasma) for free CD3 | 0.1 | 1/d | n/a |
| $k_{3,1bm}$ | Intercompartmental rate constant (plasma to tumor) for free CD3 | 0.01 | 1/d | n/a |
| BCMA parameters |  |  |  |  |
| $k_{syn,mB}$ | Synthesis rate of membrane-bound BCMA; calculated | 19.8 | nM/d | Calculated based on half-life (8) |
| $k_{shed}$ | Shedding rate of BCMA, estimated | 0.05 | 1/d | Estimated as mean of range provided in (9) |
| $k_{int\_mB}$ | Internalization rate of BCMA, calculated from half-life | 2 | 1/d | (10) |
| **Nonzero initial conditions** |  |  |  |  |
| $CD3_{th}(0)$ | Initial free CD3 target in thymus | 0.001 | nM | Estimated based on (4,7) |
| $CD3_p(0)$ | Initial free CD3 target in plasma | 0.01 | nM | |
| $CD3_{bm}(0)$ | Initial free CD3 in the bone marrow | 0.1 | nM | |
| $sB_p(0)$ | Initial free soluble BCMA in plasma | 20 | nM | Estimated based on (6) |
| $mB_{bm}(0)$ | Initial membrane-bound BCMA in the bone marrow | 50 | nM | |
| $sB_{bm}(0)$ | Initial soluble BCMA in the bone marrow | 3 | nM | |

**Table S2.** Summary of doses and near-optimal ranges that are predicted to maximize PAT (optimal dose) for the full population and for various subpopulations. Low = below median, high = above median.

| Optimal dose (mg/kg) |  | Range (within 10% of optimal) |  | Optimal dose (mg/kg) |  | Range (within 10% of optimal) |
| --- | --- | --- | --- | --- | --- | --- |
| FULL POPULATION |  |  |  |  |  |  |
| Q1W | 1.8 | 1.3-2.5 |  | Q2W | 3 | 2.5-4 |
| SUBPOPULATIONS |  |  |  |  |  |  |
| Low sBp |  |  |  |  |  |  |
| Q1W | 1.5 | 1.3-2.16 |  | Q2W | 3 | 2.16-3.5 |
| High sBp |  |  |  |  |  |  |
| Q1W | 2.5 | 1.5-2.5 |  | Q2W | 3.5 | 3-5 |
| Low sBp, Low mBCMA |  |  |  |  |  |  |
| Q1W | 1.3 | 1-1.8 |  | Q2W | 2.5 | 1.8-3 |
| Low sBp, High mBCMA |  |  |  |  |  |  |
| Q1W | 2.16 | 1.5-2.5 |  | Q2W | 3.5 | 3-5 |
| High sBp, Low mBCMA |  |  |  |  |  |  |
| Q1W | 1.5 | 1.3-2 |  | Q2W | 3 | 2.16-3.5 |
| High sBp, High mBCMA |  |  |  |  |  |  |
| Q1W | 2.5 | 1.8-3 |  | Q2W | 4.5 | 3-6 |

